# NINJ1 is a restriction factor for HSV-1 in mouse macrophages

**DOI:** 10.1101/2024.10.29.620830

**Authors:** Ella Hartenian, Magalie Agustoni, Petr Broz

## Abstract

Restriction factors block multiple stages of viral infection. Here we describe how NINJ1 controls HSV-1 infection of mouse macrophages, a key cell type during systemic and local acute infection. We observed that *Ninj1*^−/–^ macrophages are more susceptible to infection than WT cells. *Ninj1*^−/–^ macrophages exhibited earlier and stronger expression of all kinetic classes of viral proteins, resulting in higher production of infectious viral particles, with initial differences seen in the first few hours post infection. Given the important role of NINJ1 during cell death, we next investigated if NINJ1 restricts HSV-1 in a manner related to its role in cell death. However, infected *Ninj1*-deficient cells did not exhibit differences in cell death compared to WT controls at early time points in infection where we observe a difference in infection rates. Instead, the higher infection rate in *Ninj1*^−/–^ macrophages correlated to enhancement in the average numbers of viral particles entering each cell. Finally, we determined the consequences of the altered entry and found higher ISG RNA expression in infected *Ninj1*^−/–^ cells, which we ascribe to both TLR signaling and STING-mediated recognition. Together, this indicates that the NINJ1 present on WT macrophages reduces HSV-1 entry, which has consequences for the inflammatory phenotype associated with HSV-1 infection.

## Introduction

Herpes simplex virus 1 (HSV-1) affects 1 in 5 people worldwide, making it one of the most ubiquitous viruses. Infections are life-long, and symptoms vary between individuals, ranging from asymptomatic to blisters and ulcers, and even to severe outcomes including blindness and encephalitis. Severe symptoms are more likely to occur in immunocompromised individuals and work elucidating genetic susceptibilities to infection has provided mechanistic insight into virus-host interactions. HSV-1 initially infects keratinocytes and fibroblasts, making its way to axons which then act as a conduit for HSV-1 to attain access to the brain. Once inside the brain, HSV-1 establishes latency in sensory neurons and undergoes rounds of reactivation. When infection spreads beyond sensory neurons to other parts of the central nervous system (CNS), patients experience encephalitis – brain swelling and potentially debilitating symptoms.

Macrophages are innate immune sentinels and are competent for the expression of many interferon stimulated genes (ISGs), pattern recognition receptors (PRRs) and cell death pathways. While for many years keratinocytes, fibroblasts and neurons were considered the most relevant cell types for Herpes Simplex Virus 1 (HSV-1) infections, recent work has highlighted a key contribution of macrophages and monocyte-like cells. Indeed, a recent single cell analysis of an encephalitis model found that monocyte-like cells represented more than half of infected cells in the brain(1). Monocyte-like cells infiltrated the brain 4 days post infection and peaked at 7 days post infection. A second study found that depleting macrophages from mice during a systemic i.v. infection or a vaginal infection resulted in decreased early control of the virus in the systemic case and death of the animals in both models, highlighting the role of these important immune cells (2). Thus, how HSV-1 infects macrophages and balances infection with defense has important consequences for the outcome of infection.

One mechanism that host cells employ to limit viral replication is the induction of programmed cell death pathways, including pyroptosis, apoptosis and necroptosis. These pathways can act as sensors for viral molecules or virus induced disruption of cellular homeostasis. Suppression of apoptosis by viruses on the other hand, engages necroptosis, which serves as a back-up pathway of cell death in case apoptosis is inhibited (6). Previous reports have shown that HSV-1 modulates multiple cell death pathways during infection, including necroptosis, apoptosis and pyroptosis, inhibiting caspase-8, AIM2 and NLRP1-driven detection (7–11).

Here we describe a role for Ninjurin-1 (NINJ1), a plasma membrane resident protein highly expressed in the monocyte lineage of both human and mouse, in the restriction of HSV-1 infections. NINJ1 was originally described to be important for cell-to-cell adhesion (12), but has recently been shown to be essential for cellular lysis, the terminal event of multiple cell death pathways including apoptosis, ferroptosis, pyroptosis and necroptosis (13, 14). NINJ1 undergoes self-oligomerization during cell death to form amphipathic filaments. These filaments open during the final stages of cell death, creating large lesions in the plasma membrane that allow for the release of large cellular molecules such as danger associated molecular patterns (DAMPs), chemokines and cytokines (15). Most recently, NINJ1 has been suggested to be a sensor of mechanical membrane stress, such that NINJ1 in the plasma membrane represents predetermined “weak links” upon additional stress (16). NINJ1 has further been shown to be important for the release of norovirus, a non-enveloped virus that both triggers and requires cell death for egress (17).

Here we describe how infection with *Ninj1*-deficient mouse macrophages leads to greater infection of HSV-1. By monitoring HSV-1 GFP infection dynamics we see earlier and stronger infection of *Ninj1*^*–/–*^ macrophages as compared to WT controls. This affects all stages of the HSV-1 lifecycle and results in higher production of infectious viral particles, although we observe initial differences shortly after infection. We next investigate if NINJ1 restricts HSV-1 in a manner related to its role in cell death. However, we observed that infected *Ninj1*-deficient cells do not die more or earlier than WT controls at early time points in infection where we observe a difference in infection rates. We do, on the other hand, observe a higher infection rate of *Ninj1*^*–/–*^ macrophages with more viral particles entering each cell on average. Finally, we determine the consequences of the altered entry, measuring higher ISG RNA expression in infected *Ninj1*^−/–^ cells which we ascribe to both TLR signaling and STING-mediated recognition. Together this indicates that the NINJ1 present on WT macrophages reduces HSV-1 entry, which reduces the inflammatory potential of macrophages.

## Results

### NINJ1 controls susceptibility of macrophages to HSV-1

NINJ1 is a membrane protein known to have roles in cell adhesion and cell death. As both of these processes are important for pathogens to successfully invade and replicate in host cells, we sought to determine if NINJ1 plays a role during infection with HSV-1. We infected bone marrow derived macrophages (BMDMs) from WT mice or mice deficient in *Ninj1*^*–/–*^ with HSV-1 expressing a green fluorescence protein (GFP) under the control of the immediate early protein promoter ICP47. Measuring the percentage of GFP-positive (GFP+) cells as a readout for viral susceptibility, we found that *Ninj1*-deficiency resulted in twice as many infected cells compared to WT cells, as measured by flow cytometry at 6 hours post infection (hpi) (**Figure 1a**). Further analysis of infected cells showed that in the absence of NINJ1, not only were more cells infected (e.g. GFP+), they had a higher mean fluorescent intensity (MFI), indicating that NINJ1 might suppress the replication of HSV-1 in macrophages (**Figure 1b**). Similarly, widefield microscopy showed a striking increase in GFP+ cells in the absence of NINJ1 (**Figure 1c**). To understand the kinetics of HSV-1 control by NINJ, we analyzed the GFP signal intensity of infected cells every 10 minutes from the time of infection to 15 hpi. We observed that after an initial drop in the GFP signal, which occurred in both genotypes, there was a stronger increase in the signal in *Ninj1*^*–/–*^ cells as compared to the WT cells, starting around 4 hours (h) and plateauing around 11 h (**Figure 1d**), consistent with our other measures of HSV-1 GFP signal.

**Figure 1.**
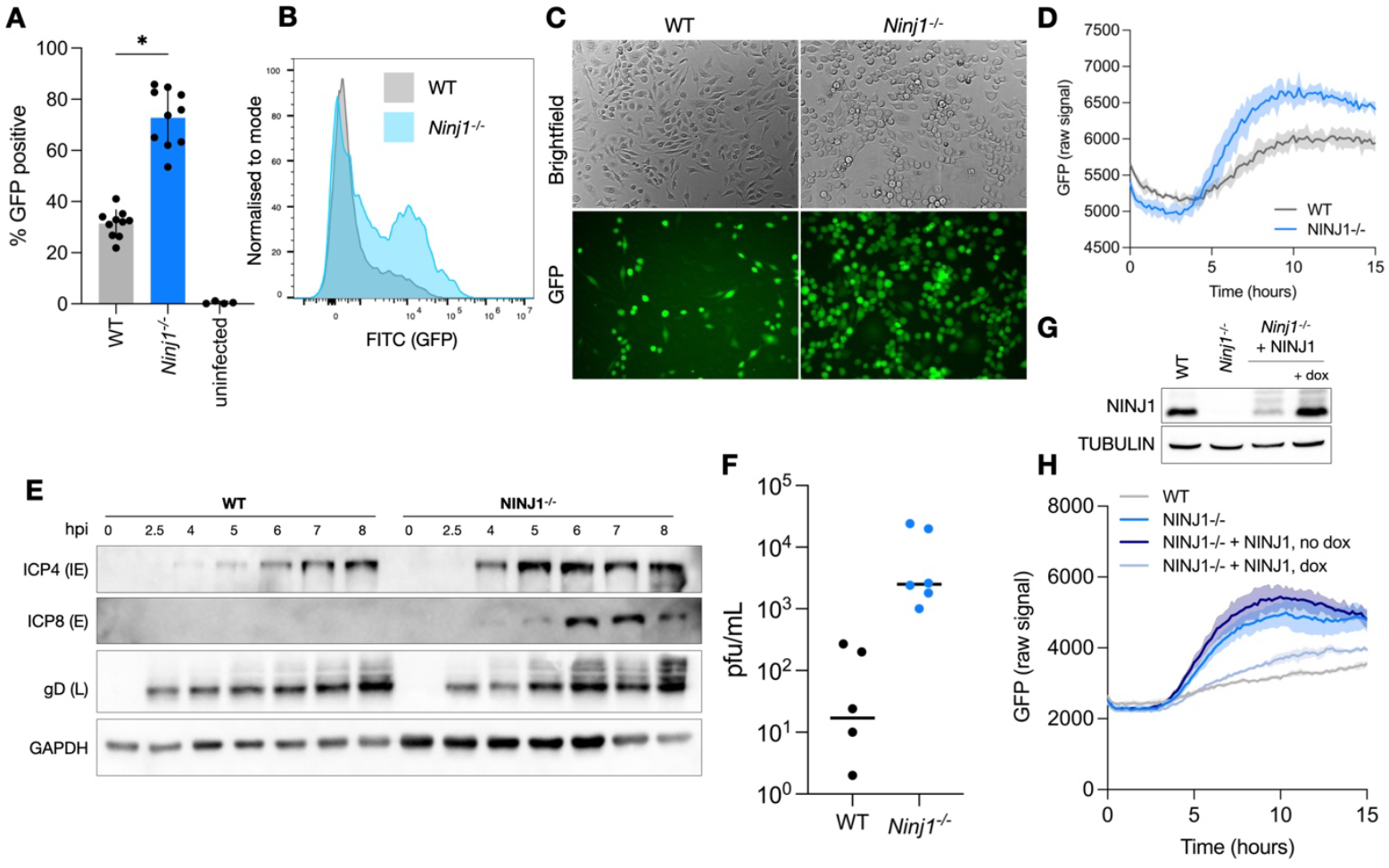
NINJ1 controls the susceptibility of mouse macrophages to HSV-1. **A-B** WT and *Ninj1*^*–/–*^ iBMDMs were infected with HSV-1 GFP for 6 h at a MOI of 10 and (A) subjected to flow cytometry to determine the percentage of infected cells, (B) the MFI. **C** WT and *Ninj1*^*–/–*^ primary BMDMs were infected and visualized by microscopy after 6 h. **D** WT and *Ninj1*^*–/–*^ iBMDMs were infected at a MOI of 10 and the GFP signal was measured every 10 min with a Citation5 Plate Reader. **E** WT and *Ninj1*^*–/–*^ iBMDMs were infected with HSV-1 GFP for the indicated timepoints then subjected to Western Blotting for the indicated proteins. **F** Plaque assays from supernatant of WT and *Ninj1*^*–/–*^ iBMDMs infected with HSV-1 GFP for 24 h. **G** Western blot analysis of NINJ1 levels in WT and *Ninj1*^*–/–*^ iBMDMs complemented with a doxycycline (dox) inducible NINJ1 in the presence and absence of dox. TUBULIN serves as a loading control. **H** GFP signal measured every 10 minutes in WT, *Ninj1*^*–/–*^ iBMDMs and complemented *Ninj1*^*–/–*^ iBMDMs +/-dox with a Citation5 Plate reader.

To confirm that the increase in HSV-1 GFP signal intensity corresponded to an increase in viral gene products, we analyzed the expression of three viral proteins from different kinetic classes – ICP4 (immediate early), ICP8, (early) and gD (late) – from 2.5 hpi to 8 hpi by immunoblotting in infected immortalized bone marrow derived macrophages (iBMDMs). Consistent with faster replication of HSV-1 in absence of NINJ1, *Ninj1*^*–/–*^ macrophages showed higher expression levels of all three proteins and earlier expression of ICP4 and ICP8 (**Figure 1e**). We next queried if the higher levels of viral protein expression we observed resulted in the production of more infectious virions in these cells. We therefore collected supernatants from cells infected for 24 h and measured the number of infectious virions by plaque assay. We observed two logs higher titer from virus that replicated in *Ninj1*^*–/–*^ cells as compared to WT cells (**Figure 1f**), demonstrating that *Ninj1*^*–/–*^ cells have faster infection kinetics and produce more virions than WT cells.

To confirm that the observed phenotype was due to *Ninj1*-deficiency alone, we transduced *Ninj1*^*–/–*^ iBMDMs with a vector expressing *Ninj1* under a doxycycline(dox)-inducible promoter. Treatment with dox induced expression of NINJ1 to levels comparable to WT cells (**Figure 1g**). A low level of background NINJ1 expression was detectable in absence of dox, most likely due to leaky expression. We next infected WT, *Ninj1*^*–/–*^ and *Ninj1*^*–/–*^ *+ Ninj1* iBMDMs, in the presence or absence of dox, with HSV-1 GFP and measured the GFP signal over time. We observed that complementing *Ninj1*-deficiency reduced the GFP signal to levels comparable to WT cells, confirming that *Ninj1*-deficiency was indeed the reason for increased viral replication (**Figure 1h**). In the complemented cells in the absence of dox, the GFP signal followed the same trend as in *Ninj1*^*–/–*^ cells, however was slightly attenuated, likely due to the leaky expression of the construct. In summary, the results show that NINJ1 controls the susceptibility of BMDMs to HSV-1.

### NINJ1 prevents infection in a cell-death independent manner

Given the key role of NINJ1 in cell lysis downstream of multiple programmed cell death (PCD) pathways and the implication of multiple of these pathways in defense against HSV-1, we wished to determine if there was redundancy between *Ninj1*-deficiency and a PCD pathway, implicating it upstream of NINJ1 activation. We therefore tested three knockout cell lines deficient for key cell death pathway components: macrophages deficient for ASC (encoded by the gene *Pycard*), a key protein for pyroptosis; *iRipk3*^*–/–*^ cells, immortalized BMDMs that lack Receptor interacting protein kinase 3 (RIPK3), a critical necroptosis adaptor protein, and are unable to undergo necroptosis; and *Casp3*^*–/–*^ BMDMs that lack the executioner caspase-3 and are thus deficient for apoptosis. We infected all three of these cell lines in parallel with WT and *Ninj1*^*–/–*^ cells with HSV-1 GFP and measured the GFP signal over time. We observed a clear increase in the GFP signal in the *Ninj1*^*–/–*^ cells, but not in any of the other cell lines, suggesting that the individual removal of each of these cell death pathways did not increase HSV-1 GFP replication and phenocopy the *Ninj1*^*–/–*^ macrophages (**Figure 2a**).

**Figure 2.**
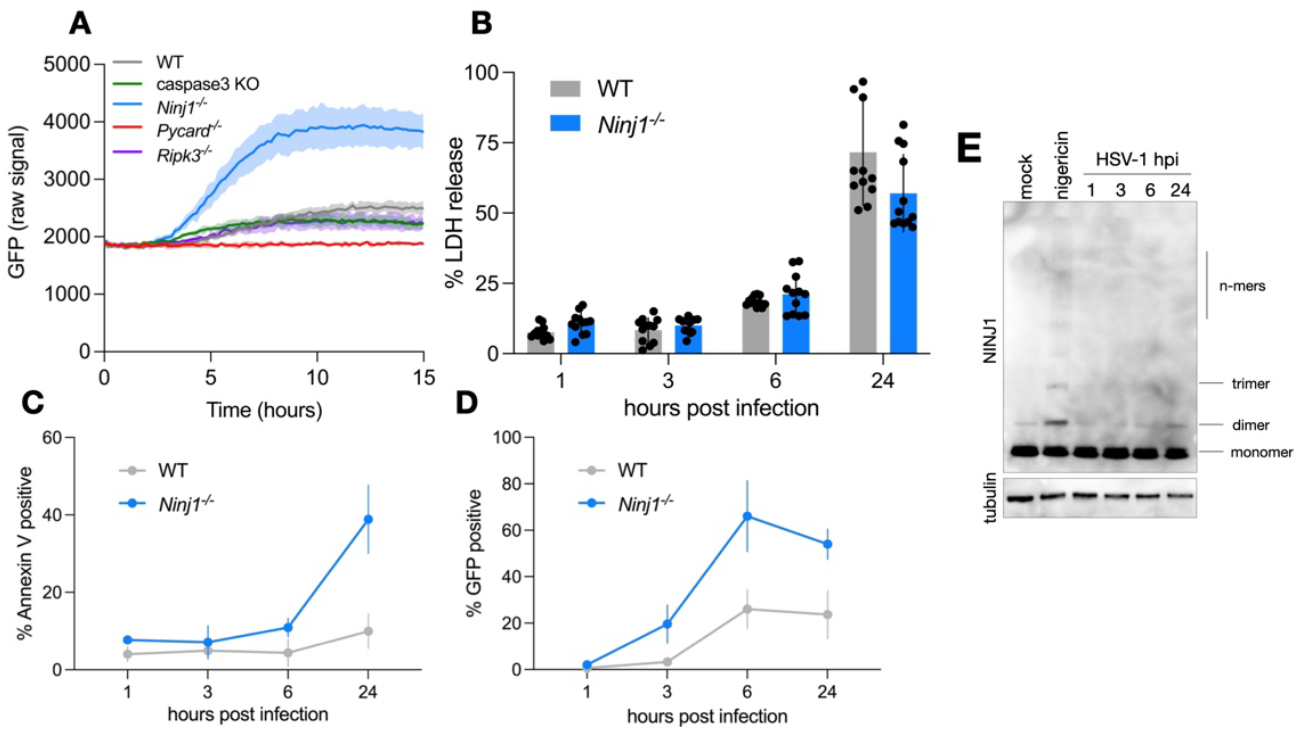
NINJ1 prevents infection in a cell-death independent manner. **A** The indicated genotypes of iBMDMs were infected with HSV-1 GFP and GFP signal was monitored every 10 minutes. **B** Lactate dehydrogenase (LDH) release was measured from the supernatant of HSV-1 GFP infected WT and *Ninj1*^*–/–*^ iBMDMs at the indicated hours post infection. **C-D** WT and *Ninj1*^*–/–*^ iBMDMs infected with HSV-1 GFP were stained with APC Annexin-V and subjected to flow cytometry, analyzing the Annexin V APC signal (C) and the percentage of infected cells (D). **E** Cross-linking immuno-blot of WT primary BMDMs infected with HSV-1 GFP for the indicated periods of time. H3 serves as a loading control and nigericin shows oligomerized NINJ1 upon cell death.

We then set out to determine if the absence of NINJ1 delayed macrophage death, thus allowing HSV-1 to replicate to higher levels. We therefore measured the release of Lactate dehydrogenase (LDH), a marker for lytic cell death, from HSV-1 infected WT and *Ninj1*^*–/–*^ cells at 1, 3, 6 and 24 hpi. Even though *Ninj1*-deficiency had an impact on viral replication as early as 5-6 hpi (**Figure 1**), we did not observe any difference in LDH release between WT and *Ninj1*^*–/–*^ macrophages at these timepoints, which remained below 25% for both cell types (**Figure 2b**). By 24 hpi, LDH release had increased in both genotypes to around 60%, but was only 10% higher in WT cells than *Ninj1*^*–/–*^ cells.

While cells were not measurably lysing, lysis is only the terminal stage of lytic cell death pathways such as pyroptosis and necroptosis, while apoptosis does not induce cell lysis. Thus cells could still be undergoing cell death which could then be incomplete or inhibited. We thus used Annexin-V staining which measures the exposure of phosphatidyl serine on the plasma membrane. Phosphatidyl serine flipping from the inner leaflet to the outer leaflet of the plasma membrane is a general hallmark of programmed cell death pathways and should therefore be agnostic to any one specific pathway. Annexin V staining showed low levels of phosphatidyl serine in both HSV-1 infected WT and *Ninj1*^*–/–*^ cells at 1-6 hpi (**Figure 2c**). At 24hpi, the Annexin V signal increased significantly in the *Ninj1*^*–/–*^ cells while remaining lower in the WT cells. The increase in the *Ninj1*^*–/–*^ cells corresponded to a higher infection rate, as measured by GFP positivity in the same samples, which peaked in the *Ninj1*^*–/–*^ cells at 6h and declined slightly by 24h (**Figure 2d**).

A hallmark of NINJ1 activation during cell death is oligomerization of the protein, which is required to induce cell lysis. Since our data suggest that NINJ1 plays a cell-lysis independent role during HSV-1 infections, we tested if NINJ1 oligomerized during HSV-1 infection by infecting cells with HSV-1 and measured NINJ1 oligomerization by adding the crosslinking agent BS3. As a control we included cell treated with nigericin, an activator of the NLRP3 inflammasome, which causes pyroptotic cell death. As expected, we found that nigericin treatment resulted in the formation of NINJ1 dimer, trimers and higher-order oligomers, indicating NINJ1 activation (**Figure 2e**). By contrast, we didn’t observe any higher order oligomers in cells infected with HSV-1. A low level of NINJ1 dimers was detectable, but this was comparable to untreated controls (**Figure 2e**). This indicates that infection of WT macrophages with HSV-1 does not lead to oligomerization at the timepoints between 1 and 24hpi that we tested, further confirming that NINJ1 must control HSV-1 infection independently of its role as an executioner of lytic cell death.

### Early stages of the viral lifecycle are impacted by the loss of NINJ1

Having ruled out an antiviral effect of NINJ1 related to its role in cell death, we next wished to distinguish at what stage of the viral lifecycle NINJ1 was inhibiting infection. Since we had observed a difference in GFP expression as early as 6 hpi (**Figure 1a,c,d**) and a difference in ICP8 expression at 4 hpi (**Figure 1e**), we speculated that NINJ1 modulated early events in the viral lifecycle. However, to nevertheless rule out that *Ninj1*-deficiency affected viral gene replication, late gene expression or subsequent events, we treated WT and *Ninj1*^*–/–*^ cells with the viral DNA replication inhibitor Phosphonoacetic acid (PAA) during HSV-1 GFP infection and measured the GFP signal. PAA did not rescue the higher infection rate of *Ninj1*^*–/–*^ cells, which we still observed in the presence or absence of NINJ1 at 18hpi (**Figure 3a**). Western blotting for viral gene products at 18hpi showed reduced expression of the late protein gD in the presence of PAA (**Figure 3b**), confirming the efficacy of PAA and indicating that NINJ1 controls a pre-DNA replication stage of the HSV-1 lifecycle.

**Figure 3.**
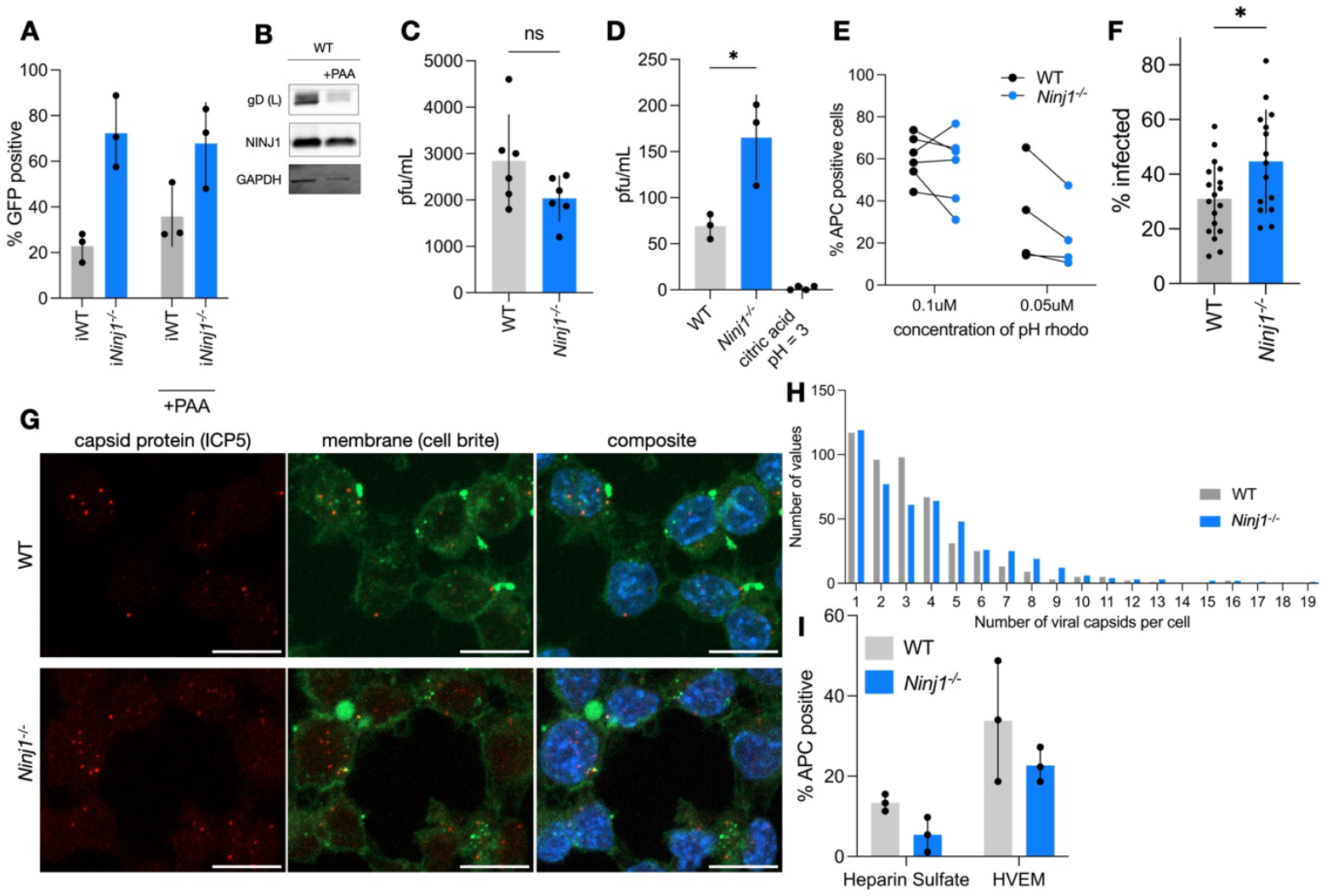
Viral entry is enhanced in the absence of NINJ1. **A** WT and *Ninj1*^*–/–*^ iBMDMs were infected for 18 h with HSV-1 GFP in the presence or absence of phosphonoacetic acid (PAA) and subjected to flow cytometry to measure the fraction of GFP positive cells. **B** Plaque assay of HSV-1 GFP incubated with WT and *Ninj1*^*–/–*^ iBMDMs for 1 h at 4°C. **C** Plaque assay of HSV-1 GFP incubated with WT and *Ninj1*^*–/–*^ iBMDMs for 15 min and then washed with low acid citric acid buffer. Citric acid condition was HSV-1 and citric acid buffer added at the same time. **D** WT and *Ninj1*^*–/–*^ iBMDMs were subjected to flow cytometry after incubation with the indicated concentration of fluorescent dextrans for 30 min. **F-H** Confocal fluorescent microscopy of cells infected with HSV-1 GFP for 45 minutes and stained with ICP5, cell brite and Hoechst. (F) Percentage of cells infected. (G) Representative images, scale bar 10um (H) Histogram of the number of capsids (ICP5 positive foci) per WT and *Ninj1*^*–/–*^ cell, defined using cellbrite stain. **I** WT and *Ninj1*^*–/–*^ iBMDMs were subjected to flow cytometry after being incubated with Heparin Sulfate and HVEM primary antibodies. F-H is from at least 20 fields of view from 3 independent experiments. F represents averages from between 750 and 950 cells.

### NINJ1 blocks viral entry into cells

Having determined that pre-DNA replication stages in the viral lifecycle are implicated in the phenotype resulting from a loss of NINJ1, we first examined viral binding, the initial step of viral entry. To measure the number of viral particles that can bind to WT and *Ninj1*^*–/–*^ cells, we incubated viral inoculum with cells at 4ºC, which allows viral binding but prevents entry, for 1 h, washed away unbound viral particles and compared the number of viral particles bound to cells by plaque assay. While we observed slightly lower pfu/mL associated with *Ninj1*^*–/–*^ cells as compared to WT (**Figure 3c**), the difference was too small to explain the large difference in HSV-1 replication between WT and *Ninj1*^*–/–*^ cells, thus suggesting that the defect in *Ninj1*^*–/–*^ cells is not a consequence of altered viral binding.

The step immediately following viral binding is viral entry. HSV-1 can enter cells both by direct membrane fusion and by endocytosis, which is then followed by the fusion of the viral particle with the endocytic membrane and the release of the capsid into the cytosol. Endocytic entry has been proposed to be the major pathway for entry into macrophages (2). Right after endocytic uptake, there is a short window of time in which viral particles remain intact and infectious within endocytic vesicles, before they fuse with the endosomal membrane. We therefore speculated that if NINJ1 restricts viral entry, more infectious virions could be recovered from endosomes in the absence of NINJ1. To assess the level of entry into cells, we thus infected WT and *Ninj1*^*–/–*^ cells for 15 min with HSV-1, treated cells with citric acid to inactivate all viral particles that had not entered cells and then determined the number of infectious virions (i.e. virions within endosomes) by plaque assay. We observed 3 times as many viral particles in *Ninj1*^*–/–*^ cells as compared to WT cells (**Figure 3d**), indicating that NINJ1 restricted HSV-1 entry into cells. To understand if this was a specific or general effect, we tested if *Ninj1-*deficiency impacted the endocytosis of fluorescent dextrans. We observed no change or even a slight decrease in endocytosis of *Ninj1*^*–/–*^ macrophages (**Figure 3e**), which could not explain our HSV-1 entry results and suggested that NINJ1 was playing a specific role in viral entry.

Since HSV-1 can also enter cells by direct fusion with the plasma membrane, we used confocal microscopy to quantify the fraction of cells positive for ICP5, a HSV-1 capsid protein after infection for 45 minutes. This method will detect virions that entered by either means, direct membrane fusion or endocytosis. We saw that twice as many *Ninj1*^*–/–*^ cells were infected as compared to WT cells (**Figure 3f,g**). We also counted the number of capsids inside each infected cell in our microscopy and observed that *Ninj1*^*–/–*^ cells tended to harbor more capsids per cell than WT cells (**Figure 3h**). This confirms that HSV-1 can both enter more cells in the absence of NINJ1 and that the number of successfully entry events per cell increases.

Finally, we asked if the absence of NINJ1 alters expression of known HSV-1 entry mediators. We compared the presence of HVEM, Heparin Sulfate and Nectin-1 on the surface of WT and *Ninj1*^*–/–*^ cells. There was slightly less Heparin Sulfate and Nectin-1 detectable on the cell surface by Fluorescence activated cell sorting (FACS) in the *Ninj1*^*–/–*^ cells as compared to WT, which could not explain higher viral entry into these cells (**Figure 3i**). Nectin-1 levels were not detectable on either genotype, with either of two different antibodies. In summary these data demonstrate that NINJ1 restricts the entry of HSV-1 into macrophages both via direct membrane fusions or via endosomes, and that this does not depend on altered expression of known HSV-1 entry mediators.

### NINJ1 is highly expressed in primary macrophages

Having characterized the role of NINJ1 as a gatekeeper for HSV-1 entry in macrophages, we wondered if this role could apply to other cell types that could be infected by HSV-1. We first looked at a panel of human cell lines to check for NINJ1 expression using an antibody we made to a N-terminal NINJ1 peptide. Comparing all cell lines to human monocyte derived macrophages (hMDMs) which have previously been shown to have robust NINJ1 expression (18), we saw very limited expression in other cell lines (**Figure 4a**). Human protein atlas RNA data similarly suggested a significantly higher amount of NINJ1 expression in bone-marrow derived cells (**Figure 4b**). Given the nearly undetectable levels of expression in common laboratory human cell lines, we therefore turned to other mouse lines, where upon blotting for NINJ1 we observed expression in mouse embryonic fibroblasts (MEFs), RAW macrophages and 3T3 fibroblasts (**Figure 4c**). To check if NINJ1 controls HSV-1 infection in these cell types as we had observed in the macrophages, we infected sgluciferase expressing control cells and CRISPR-generated NINJ1 KO cells. However, the infection of NINJ1 KO MEFs was quite similar to control cells (**Figure 4d**). In 3T3s, on the other hand, we observed slightly higher infection rates in NINJ1 KO 3T3s (**Figure 4e**), suggesting a role for NINJ1 in infection of these cells. In summary, these findings show that NINJ1 expression is mostly detectable in both primary human and mouse macrophages, and that expression of NINJ1 can be found in other cells types, including RAW macrophages and mouse fibroblasts. It is thus likely that NINJ1 is a cell type specific factor that controls plasma membrane processes, such as death-associated membrane rupture and viral entry, primarily in macrophages.

**Figure 4.**
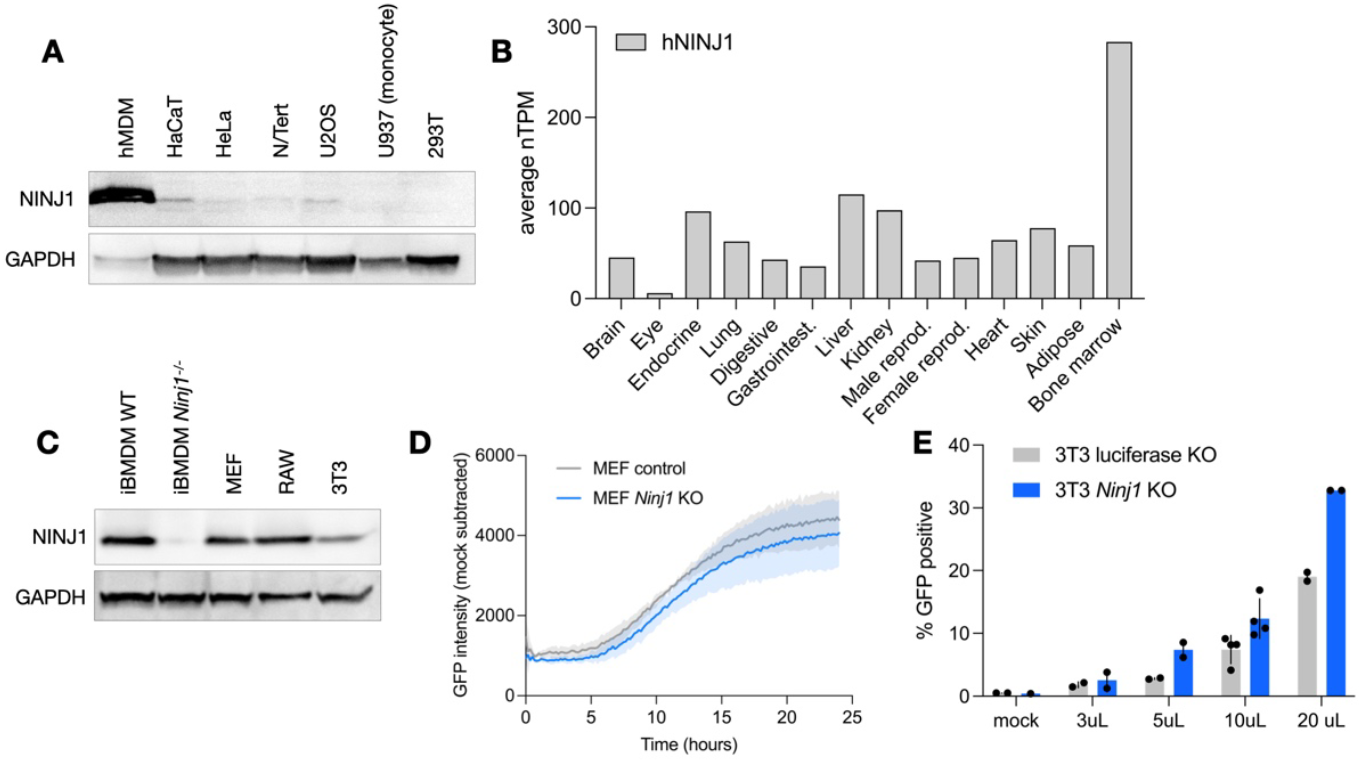
NINJ1 is highly expressed in primary macrophages. **A** The indicated cell lines were subjected to Western blotted using an antibody against human NINJ1 and GAPDH as a control. **B** Data from the human protein atlas (transcripts per million) were plotted indicating NINJ1 transcript expression in different human tissues. **C** The indicated cell lines were subjected to Western blotting using an antibody against mouse NINJ1 and GAPDH as a control. **D** Infection of control and *Ninj1* KO MEFs with HSV-1 GFP was measured by detecting the GFP signal every 10 minutes. **E** Infection of luciferase and *Ninj1* KO 3T3s was performed with different amounts of HSV-1 GFP for 8 h and infection rate was measured by flow cytometry. A and C and D are representative of 3 independent experiments.

### ISGs are upregulated in macrophages lacking NINJ1

Given our observation that more viral particles enter *Ninj1*^*–/–*^ macrophages, we finally asked how this affected the innate immune response to HSV-1. We therefore measured the levels of several ISGs in WT and *Ninj1*^*–/–*^ cells by RT-qPCR. We observed a trend of a stronger induction of ISGs in the *Ninj1*^*–/–*^ cells with the most pronounced difference in *Viperin* expression and smaller differences in *Bst1 and IFI27I2a* expression (**Figure 5a**). Interestingly, for *Viperin*, we observed a difference beginning as early as 3 hpi. We then wished to know what host pathway was driving the induction of ISGs. Both TLRs and the cGAS/STING cytosolic DNA sensing pathway have been implicated in the innate immune response to HSV-1(19–21). Since TLRs signal through the adaptor proteins MyD88 and TRIF, we checked if ISG levels were altered in *MyD88*^*–/–*^*Trif*^*–/–*^ and *Sting*^*–/–*^ iBMDMs. Comparing the induction of ISGs at 3 h and 6 h, we observed the most marked decrease in both *IFI2712a* and *Viperin* induction in *MyD88*^*–/–*^*Trif*^*–/–*^ cells (**Figure 5b**). We also observed a decrease in *Sting*^*–/–*^ cells but this was more consistent for the induction of *Viperin* as compared to the induction of *IFI2712a*. In conclusion, we implicate both TLR-mediated sensing and cytosolic DNA sensing to the macrophage response to HSV-1, which is elevated in the absence of NINJ1.

**Figure 5.**
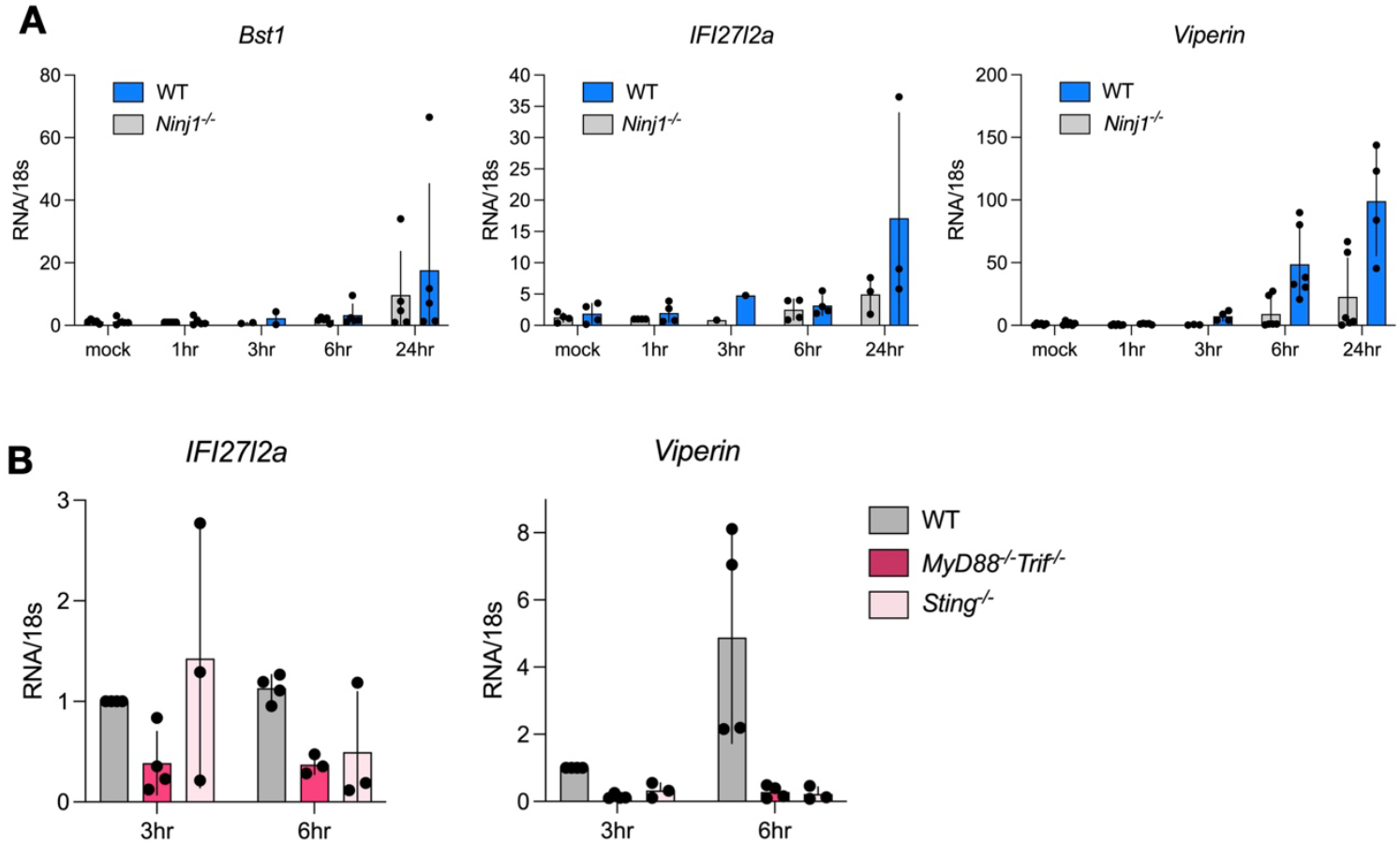
Loss of Ninj1 results in higher levels of ISGs. **A** WT and *Ninj1*^*–/–*^ iBMDMs were infected for the indicated times with HSV-1 GFP and then RNA levels of the indicated transcripts were measured by RT-qPCR. **B** WT, *MyD88*^*–/–*^*Trif*^*–/–*^ and *Sting*^*–/–*^ iBMDMs were infected with HSV-1 GFP for the indicated times and then RNA levels of the indicated transcripts were measured by RT-qPCR.

## Discussion

Here we describe how NINJ1 acts as a gatekeeper for HSV-1 in mouse macrophages. We see that the HSV-1 lifecycle proceeds more quickly in the absence of NINJ1 and results in the production of more infectious virions. We pinpoint the difference in infection rates to a difference in viral entry, with more viral particles entering *Ninj1*^*–/–*^ cells and a greater percentage of *Ninj1*^*–/–*^ cells getting infected. We can observe this both by measuring viral particles entering by endocytosis as well as by using microscopy to measure the number of capsids inside of cells. Thus, while NINJ1 has recently been identified as a key mediator of membrane rupture and cell lysis during cell death, we observe a novel and cell death independent role for NINJ1 as a plasma membrane protein that renders macrophages refractory to HSV-1 infection at the site of first virus-cell contact.

Macrophages are recruited to the site of infection during both systemic (IV) and cutaneous models. During an encephalitis model of infection, monocyte-like cells represented more than half of the total population harboring viral transcripts, indicating that HSV-1 enters these cells and completes some of the viral lifecycle (1). Mouse models of HSV-1 infection that lack macrophages succumb to disease much more quickly than controls, highlighting the key role macrophages play in controlling lytic infection (2). NINJ1 is highly expressed in macrophages in comparison to most other cell types, thus modulating the infection rate of a key cell type. Here we observe that in the absence of NINJ1, ISG expression increases, suggesting a differential host response that could be too robust if NINJ1 was absent, having consequences for tissue damage and auto-inflammation. We further define that macrophages detect HSV-1 both through TLR and cytosolic nucleic acid-dependent pathways, as removal of either of these pathways dampens the ISG response of infected cells.

In addition to being present on the plasma membrane, NINJ1 is also found on the nuclear membrane and in the secretory system (likely during trafficking to the plasma membrane)(15). HSV-1 interfaces both with the nucleus and the secretory pathway, thus while we ascribe the entry-related role of NINJ1 to the plasma membrane resident population, we cannot rule out secondary roles during infection of NINJ1 elsewhere in the cell. Furthermore, we do not observe oligomerization of NINJ1 during HSV-1 infection even though we observe macrophages releasing LDH at late stages of infection, indicative of lytic cell death. While HSV-1 is known to be able to trigger apoptosis, necroptosis and pyroptosis (8, 10, 11) HSV-1 may directly block NINJ1 oligomerization. This could happen either by a direct block of NINJ1 or by indirectly by inhibiting cell death pathways that could trigger NINJ1. Alternatively, LDH release could occur in a NINJ1-independent manner at late stages of infection.

In conclusion, here we demonstrate a novel role for NINJ1 in preventing entry of HSV-1 in mouse macrophages, a cell type that expresses the greatest amount of NINJ1 in the body and is essential for HSV-1 infection.

## Materials and Methods

### Mammalian cell culture and generation

*WT* and *Ninj1*^*–/–*^ mouse bone marrow-derived macrophages (BMDMs) were harvested and differentiated in Dulbecco’s modified Eagle Medium (DMEM) (Gibco) containing 20% L929L supernatant as a source of macrophage colony-stimulating factor (M-CSF), 10% heat-inactivated fetal calf serum (FCS) (BioConcept), 10 mM Hepes (BioConcept), 1% penicillin/streptomycin and 5 mL nonessential amino acids (NEAA, Gibco). Experiments were performed on days 9 to 10 of differentiation. Immortalization of macrophages was performed as previously described (22). Briefly, primary monocytes were extracted from bone marrow and then treated with filtered supernatant from existing iBMDMs for 24 h. Then these cells are grown for up to 2 months to wait for those that are immortalized to grow up. After several weeks in culture, M-CSF in the media is reduced from 20% to 10%. Immortalized macrophages (iBMDMs) were cultured in DMEM containing 10% FCS, 10% M-CSF, 10 mM Hepes and 5mL nonessential amino acids. HEK 293T and 3T3 cells were cultured in DMEM supplemented with 10% FCS. 3T3 KO and controls have been previously described (14). MEFs were grown in DMEM with 10% FCS, 2 mM L-glutamine, and 400 µM sodium pyruvate and were previously published (23). All cells were grown at 37°C, 5% CO_2_.

### Viral propagation, infection and plaque assays

KOS HSV-1 GFP (ICP47 promoter drives GFP, a kind gift of Florian Schmidt) was propagated on Vero cells. Briefly, 90% confluent cells were infected at an MOI of 0.01 and left to grow for 3-4 days. Cell free viral particles were combined with cells subjected to douncing, ultracentrifuged at 15 000 x g for 90 minutes and the viral pellet was resuspended in DMEM with 10% FCS and 1g/100mL of BSA. Viral preps were titered on Vero cells and MOIs reflect Vero cell numbers. Plaque assays were performed by seeding Vero cells at 1 × 10^4^ cells per well in 24-well plates. The next day cells were infected in 500uL for 2 h, then aspirated and replaced with full media supplemented with 0.5% carboxymethylcellulose, then left to grow for 3 days. Cells were fixed with 4% paraformaldehyde for 10 minutes, then aspirated and stained with 0.5% crystal violet for 10 minutes before counting.

Infections were performed at an MOI of 10 when not indicated otherwise. For macrophage infections, viral supernatants were left on cells for 3 h in optiMEM before changing media, except for experiments analyzed by BioTek Cytation5 plate reader where virus was added to cells and not removed.

### Immunoblotting

For western blotting analysis, cells were lysed in 66 mM Tris-HCl pH 7.4, 2% SDS, 10 mM DTT, and NuPage LDS sample buffer (Thermo Fisher Proteins were separated on gradient precast gels 4-20% (Milipore) and transferred onto nitrocellulose membrane using Transblot Turbo (Bio-Rad). The antibodies used were: anti-ICP4 (Santa Cruz 56986), anti-ICP8 (Abcam 20194), anti-gD (Santa Cruz 69802) anti-mouse NINJ1 (rabbit IgG2b clone 25; a kind gift from Genentech; 1:8000), anti-tubulin (ab40742; Abcam; 1:2000), anti-GAPDH (365062; Santa cruz, 1:3000). Primary antibodies were detected with horseradish peroxidase (HRP)-conjugated goat anti-rabbit (4030-05; Southern Biotech; 1:5000), HRP-conjugated goat anti-mouse (1034-05; Southern Biotech; 1:5000 or 12-349; MilliporeSigma; 1:2000) secondary antibodies.

### Complementation of NINJ1 in NINJ1–/– iBMDMs

pLVX mNINJ1 (addgene 208775) was made into lentivirus by transfecting 3 million 293T cells in a 6-well dish with 1.25*µ*g of packaging plasmid (pLVX mNINJ1), 1.25µg of psPAX2 and 250ng of VSVG and 8uL LT-1 transfection reagent (Mirus). Media was changed 24 h post transfection to DMEM + 10% FCS + 1g/100mL Bovine Serum Albumin (BSA), lentivirus was harvested 48 h post transfection and filtered through a 0.45 µm filter. 1.5 million iBMDMs were transduced with 1mL of virus by spinning at 500xg for 1.5 h in one well of a non tissue culture treated 6-well dish. Next, cells were split out into a 10cm petri dish and selected with 1µg/mL blasticidin 24 h post transduction for at least 5 days.

### Entry experiments

For infections where entry by endocytosis was quantified, after addition of viral supernatants onto cells, cells were washed with a low pH citric acid buffer (135mM NaCl, 10mM KcL, 40mM citric acid, pH 3) for 5 min to inactivate any extracellular viral particles.

### Annexin V flow cytometry

Macrophages were seeded in tissue culture treated 12-well plates at 1×10^5^ cells / well. Supernatant and adherent cells were combined by trypsinzing adherent cells and then stained with Annexin V APC (BioLegend) following the manufacturer’s protocol, using BioLegend Cell Staining Buffer and Annexin V Binding Buffer (BioLegend)

### Cell lysis assays

A day before stimulation, cells were seeded in 96-well plates at a density of 2 × 10^4^ iBMDMs. Cells were infected at an MOI of 10. Cell lysis was quantified by measuring the lactate dehydrogenase (LDH) amount in the cell supernatant using the LDH cytotoxicity kit (Takara, Clontech) according to the manufacturer’s instructions and expressed as a percentage of total LDH release. LDH release was normalized to untreated control and 100% lysis control, by adding Triton X-100 to a final concentration of 0.01%: (LDH_sample_ – LDH_negative control_)/(LDH_100% lysis_ – LDHnegative control) x 100.

### Crosslinking assay

WT BMDMs were seeded in 24-well plates at a density of 2 × 10^5^ cells per well a day before stimulation. The next day, cells were infected with HSV-1 at MOI of 50 for 3 h, or the relevant time post infection. Cells were washed once in Ca^2+^Mg^2+^ containing PBS and then 200uL of Ca^2+^Mg^2+^containing PBS was left on the cells. Next, the crosslinker BS^3^ (bis(sulfosuccinimidyl)suberate) was added according to the manufacturer’s instructions. In brief, BS^3^ was added to the media (3 mM; Thermo Fisher) and incubated for 5 min at room temperature. Next, a solution of 20mM Tris pH 7.5 was added to stop the reaction and incubated for 10 min at room temperature. Cell supernatants were collected, and proteins were precipitated with methanol and chloroform and combined with cell lysates for western blotting analysis.

### Confocal microscopy and image analysis

iBMDMs seeded onto glass coverslips were infected for 45minutes at a MOI of 10. For the last 5 minutes of infection they were treated with CellBrite (Biotium) to label the plasma membrane. Then cells were fixed in 4% PFA for 10 minutes, washed with PBS, permeabilized with 0.05% saponin and blocked with 1% BSA in PBS. Then, samples were incubated with an anti-ICP5 antibody (abcam 6508), then an Alexa Fluor-568 conjugated antibody and Hoechst (1:1000). Samples were then imaged with a Zeiss LSM800 confocal laser scanning microscope using a 63x/1.4 NA oil objective. Quantification of capsids was performed using a maximum projection of the z-stack, by manually segmenting cells and counting the number of infected cells per field of view and the number of capsids per cell. All microscopy datasets were analyzed and processed using Fiji software.

### Dextran uptake assays

iBMDMs were seeded at 5×10^5^ in 100uL of FACS buffer (PBS with xx) in a V bottom 96-well plate. Dextran was added to a final concentration of either 0.05uM or 0.1uM and incubated at 37°C for 30 min with gentle agitation every 10 min. Each well was then supplemented with 100uL FACS buffer, spun at 300xg for 5m at 4°C, washed once with 200uL FACS buffer, spun again as before and resuspended in 250uL FACS buffer for analysis. After the initial 30 min incubation samples were kept on ice to prevent additional Dextran entry until being analyzed by flow cytometry.

### Antibody generation

Rabbit anti human NINJ1 was made by YenZym antibodies, LLC to a synthetic NINJ1 peptide comprising the first 76 aa of human NINJ1 (GenScript).

### RT-qPCR

RNA was isolated with Trizol. Briefly, 1×10^5^ cells seeded the day before and then infected were washed and resuspended in 50uL of PBS, then lysed in 500uL of Trizol. 100uL of chloroform was added and spun for 15m at 21,000 x g at 4C. The top phase was moved into a new tube with 5uL of glycogen and 500uL of isopropanol. After 10 min at room temperature, samples were spun again for 10m at 21,000 x g at 4C, and then washed twice with 70% Ethanol, spinning for 5 min after each spin. Then Ethanol was removed, samples were resuspended in Rnase/Dnase free water. 1.5ug of RNA was Dnase treated and then subjected to RT with MMLV. RT-qPCR was performed with SyberGreen Master Mix.

### Animals and primary cells

All experiments implicating animals were performed under the guidelines and approval from the cantonal veterinary office of the canton of Vaud (Switzerland), license number VD3895. All mice were bred and housed at a specific-pathogen-free facility at 22 C° room temperature, 10 % humidity and a day/night cycle of 12h/12h at the University of Lausanne. *Ninj1*-deficient mice have been described before (Degen et al). *Myd88*^*–/–*^*Sting*^*–/–*^, *Sting*^*–/–*^, *Ripk3*^−/–^, *Pycard*^−/–^ (ASC) and caspase 3 KO cells have been described before (24, 25).

## Acknowledgments

This work is supported by ERC-CoG 770988 (InflamCellDeath) and SNF Project funding (310030B_198005, 310030B_192523) to P.B., EMBO postdoctoral fellowships ALTF 27-2022 to E.H. We would like to thank the UNIL animal facility for their help and support, Florian Schmidt for his kind gift of HSV-1 GFP, Mathieu Bertrand (VIB) for his gift of control and NINJ1 KO MEFs, and the Broz lab for helpful discussions.

## Disclosure and competing interest statement

The authors declare that they have no conflict of interest.

## Notes

### Competing Interest Statement

The authors have declared no competing interest.

## References

1. Ding, X., Lai, X., Klæstrup, I.H., Jensen, S.R.N., Nielsen, M.M., Thorsen, K., Romero-Ramos, M., Luo, Y., Lin, L., Reinert, L.S., et al. (2024) Temporally resolved single-cell RNA sequencing reveals protective and pathological responses during herpes simplex virus 1 CNS infection. bioRxiv, 10.1101/2024.08.19.608535.

2. Lang, J., Bohn, P., Bhat, H., Jastrow, H., Walkenfort, B., Cansiz, F., Fink, J., Bauer, M., Olszewski, D., Ramos-Nascimento, A., et al. (2020) Acid ceramidase of macrophages traps herpes simplex virus in multivesicular bodies and protects from severe disease. Nat. Commun., 11, 1338.

3. Döhner, K., Cornelius, A., Serrero, M.C. and Sodeik, B. (2021) The journey of herpesvirus capsids and genomes to the host cell nucleus. Curr. Opin. Virol., 50, 147–158.

4. Broz, P. and Dixit, V.M. (2016) Inflammasomes: mechanism of assembly, regulation and signalling. Nat Rev Immunol, 16, 407–420.

5. Mocarski, E.S., Upton, J.W. and Kaiser, W.J. (2012) Viral infection and the evolution of caspase 8-regulated apoptotic and necrotic death pathways. Nat Rev Immunol, 12, 79–88.

6. Kaiser, W.J., Upton, J.W. and Mocarski, E.S. (2013) Viral modulation of programmed necrosis. Curr Opin Virol, 3, 296–306.

7. Guo, H., Gilley, R.P., Fisher, A., Lane, R., Landsteiner, V.J., Ragan, K.B., Dovey, C.M., Carette, J.E., Upton, J.W., Mocarski, E.S., et al. (2018) Species-independent contribution of ZBP1/DAI/DLM-1-triggered necroptosis in host defense against HSV1. Cell Death Dis., 9, 816.

8. Guo, H., Koehler, H.S., Mocarski, E.S. and Dix, R.D. (2022) RIPK3 and caspase 8 collaborate to limit herpes simplex encephalitis. PLoS Pathog., 18, e1010857.

9. Maruzuru, Y., Ichinohe, T., Sato, R., Miyake, K., Okano, T., Suzuki, T., Koshiba, T., Koyanagi, N., Tsuda, S., Watanabe, M., et al. (2018) Herpes Simplex Virus 1 VP22 Inhibits AIM2-Dependent Inflammasome Activation to Enable Efficient Viral Replication. Cell Host Microbe, 23, 254–265.e7.

10. Parameswaran, P., Payne, L., Powers, J., Rashighi, M. and Orzalli, M.H. (2024) A viral E3 ubiquitin ligase produced by herpes simplex virus 1 inhibits the NLRP1 inflammasome. J. Exp. Med., 221, e20231518.

11. Aubert, M. and Blaho, J.A. (2001) Modulation of apoptosis during herpes simplex virus infection in human cells. Microbes Infect., 3, 859–866.

12. Lee, H.-J., Ahn, B.J., Shin, M.W., Choi, J.-H. and Kim, K.-W. (2010) Ninjurin1: a potential adhesion molecule and its role in inflammation and tissue remodeling. Mol. Cells, 29, 223–227.

13. Kayagaki, N., Kornfeld, O.S., Lee, B.L., Stowe, I.B., O’Rourke, K., Li, Q., Sandoval, W., Yan, D., Kang, J., Xu, M., et al. (2021) NINJ1 mediates plasma membrane rupture during lytic cell death. Nature, 591, 131–136.

14. Ramos, S., Hartenian, E., Santos, J.C., Walch, P. and Broz, P. (2024) NINJ1 induces plasma membrane rupture and release of damage-associated molecular pattern molecules during ferroptosis. EMBO J., 43, 1164–1186.

15. Degen, M., Santos, J.C., Pluhackova, K., Cebrero, G., Ramos, S., Jankevicius, G., Hartenian, E., Guillerm, U., Mari, S.A., Kohl, B., et al. (2023) Structural basis of NINJ1-mediated plasma membrane rupture in cell death. Nature, 618, 1065–1071.

16. Xu, J., Zhu, Y., Xiao, F., Wang, Y., Wang, Y., Li, J., Zhong, D., Huang, Z., Yu, M., Wang, Z., et al. (2024) NINJ1 regulates plasma membrane fragility under mechanical tension. 10.21203/rs.3.rs-5237916/v1.

17. Wang, G., Zhang, D., Orchard, R.C., Hancks, D.C. and Reese, T.A. (2023) Norovirus MLKL-like protein initiates cell death to induce viral egress. Nature, 10.1038/s41586-023-05851-w.

18. Borges, J.P., Sætra, R.S., Volchuk, A., Bugge, M., Devant, P., Sporsheim, B., Kilburn, B.R., Evavold, C.L., Kagan, J.C., Goldenberg, N.M., et al. (2022) Glycine inhibits NINJ1 membrane clustering to suppress plasma membrane rupture in cell death. Elife, 11, e78609.

19. Lima, G.K., Zolini, G.P., Mansur, D.S., Lima, B.H.F., Wischhoff, U., Astigarraga, R.G., Dias, M.F., Silva, M. das G.A., Béla, S.R., Antonelli, L.R. do V., et al. (2010) Toll-Like Receptor (TLR) 2 and TLR9 Expressed in Trigeminal Ganglia are Critical to Viral Control During Herpes Simplex Virus 1 Infection. Am. J. Pathol., 177, 2433–2445.

20. Zhang, S.-Y., Herman, M., Ciancanelli, M.J., Diego, R.P. de, Sancho-Shimizu, V., Abel, L. and Casanova, J.-L. (2013) TLR3 immunity to infection in mice and humans. Curr. Opin. Immunol., 25, 19–33.

21. Reinert, L.S., Lopušná, K., Winther, H., Sun, C., Thomsen, M.K., Nandakumar, R., Mogensen, T.H., Meyer, M., Vægter, C., Nyengaard, J.R., et al. (2016) Sensing of HSV-1 by the cGAS–STING pathway in microglia orchestrates antiviral defence in the CNS. Nat Commun, 7, 13348.

22. Broz, P., Moltke, J. von, Jones, J.W., Vance, R.E. and Monack, D.M. (2010) Differential Requirement for Caspase-1 Autoproteolysis in Pathogen-Induced Cell Death and Cytokine Processing. Cell Host Microbe, 8, 471–483.

23. Dondelinger, Y., Priem, D., Huyghe, J., Delanghe, T., Vandenabeele, P. and Bertrand, M.J.M. (2023) NINJ1 is activated by cell swelling to regulate plasma membrane permeabilization during regulated necrosis. Cell Death Dis., 14, 755.

24. Heilig, R., Dilucca, M., Boucher, D., Chen, K.W., Hancz, D., Demarco, B., Shkarina, K. and Broz, P. (2020) Caspase-1 cleaves Bid to release mitochondrial SMAC and drive secondary necrosis in the absence of GSDMD. Life Sci. Alliance, 3, e202000735.

25. Mariathasan, S., Newton, K., Monack, D.M., Vucic, D., French, D.M., Lee, W.P., Roose-Girma, M., Erickson, S. and Dixit, V.M. (2004) Differential activation of the inflammasome by caspase-1 adaptors ASC and Ipaf. Nature, 430, 213–218.

